# Higher adaxial stomatal density is associated with lower grain yield in spring wheat

**DOI:** 10.1101/2025.01.07.631641

**Authors:** Kajal Samantara, Elena Ivandi, Ingmar Tulva, Pirko Jalakas, Banafsheh Khalegh Doust, Anne Ingver, Merko Kärp, Gintaras Brazauskas, Mara Bleidere, Ilmar Tamm, Hanna Hõrak, Ebe Merilo

**Author notes:** shared correspondence.

## Abstract

In land plants, stomatal pores on leaf surfaces developed to control gas exchange between leaf and surrounding air, but also to enable nutrient uptake and leaf cooling. Traits such as stomatal density (SD), guard cell size, stomatal distribution between the upper (adaxial) and lower (abaxial) leaf surfaces (stomatal ratio), and stomatal aperture width exhibit notable variation across different genotypes and environments. These traits influence leaf photosynthesis, water loss, growth, productivity, and pathogen susceptibility. Here, we studied different stomatal traits of spring wheat flag leaves and their relationship with grain yield in field experiments during 2022-2023. Significant genotypic variation among adaxial and abaxial SDs and stomatal ratios was detected, whereas stomatal conductance was mostly affected by annual differences in weather. A strong negative relationship between adaxial stomatal density and grain yield was detected under all conditions, when abiotic factors (water stress or nutrient limitation) resulted in yield losses, whereas under favourable conditions, there was no significant relationship between adaxial stomatal density and grain yield. The effects of leaf surface-specific traits on yield are often overlooked in physiological and breeding experiments. Our results indicate that higher-than-optimal adaxial SD values may result in wheat yield losses under stresses imposed by future climate conditions.

## Main text

Stomata are microscopic pores, each formed by two guard cells, on the leaf surface that enable gas exchange between the leaf and the atmosphere. Stomatal traits e.g, stomatal density (SD, number of stomata per unit leaf area), stomatal distribution between leaf sides, guard cell length (GCL), and stomatal aperture width vary considerably among different species, genotypes and environmental conditions, affecting leaf and plant gas exchange (Salisbury, 1928; Maruyama & Tajima, 1990; Weyers & Lawson, 1997; James & Bell, 2000; Hetherington & Woodward, 2003; Muir, 2015; Stevens *et al*., 2021; Wall *et al*., 2023; Liu *et al*., 2023). Stomatal conductance (g_s_) that determines the rate of water vapour diffusion from leaves through stomata, is controlled by SD and stomatal aperture width. High values of g_s_ are associated with efficient photosynthesis, active water movement through the plant, and canopy cooling, which in turn result in higher yields, but also in higher water loss as a drawback (Wong *et al*., 1979; Roche, 2015). Changes in SD can affect photosynthesis, growth, stress tolerance, and production in different ways (Lawson & Matthews, 2020; Pflüger *et al*., 2024). Recent studies show that lower SD of genetically modified plants in Arabidopsis (Hepworth *et al*., 2015), barley (Hughes *et al*., 2017), wheat (Dunn *et al*., 2019), and rice (Caine *et al*., 2019) led to improved drought tolerance without negatively affecting growth or carbon assimilation, whereas high SD was related with hampered growth in Arabidopsis (Tulva *et al*., 2024; Jalakas *et al*., 2024).

Depending on plant species and environment, stomata can be present on only one leaf surface or on both; amphistomatous species with stomata on both leaf surfaces have different stomatal ratios (SR, ratio of adaxial SD to abaxial SD; Salisbury, 1928; Mott *et al*., 1982; Muir, 2015). Most amphistomatous species have higher SD on the abaxial leaf surface (Pemadasa, 1979; Muir, 2015; Caseys *et al*., 2024), whereas crops of the *Gramineae* family (like wheat) may have equal or even higher SD on the adaxial surface (Pathare *et al*., 2020; McAusland *et al*., 2021; Wall *et al*., 2022, 2023). The presence of stomata on both leaf sides results in enhanced photosynthetic rates via boosting stomatal and mesophyll conductances for CO_2_ diffusion, with potential disadvantages such as pathogen sensitivity or higher dehydration risk (Parkhurst, 1978; Drake *et al*., 2019; Muir, 2020; Pathare *et al*., 2020; Xiong & Flexas, 2020). Recently, it was found that adaxial and abaxial leaf surfaces mount different chemical defences, and adaxial SD was not correlated with sensitivity to *Botrytis cinerea* (Caseys *et al*., 2024). This suggests that adaxial stomata do not necessarily increase susceptibility to pathogens. Higher dehydration risk associated with adaxial stomata may be due to longer distance to the closest veins and larger leaf-to-air vapour pressure differences representing higher evaporative demand (Drake *et al*., 2019).

Globally, wheat production is reaching 800 Mt; wheat is the second most produced cereal worldwide, accounting for about one-fifth of global human caloric intake (Shiferaw *et al*., 2013; FAOSTAT, 2022). European wheat yields have stagnated since the early 1990s due to climate trends, but also changes in agricultural and environmental policies (Moore & Lobell, 2015; Le Gouis *et al*., 2020). This has led to discussions about whether and how much it is possible to boost crop yields with photosynthetic improvements (Faralli & Lawson, 2020; Matthews & Burgess, 2024) and breeding plants with higher production in well-watered *versus* drought-prone environments by modifying SD (Hepworth *et al*., 2018).

Here, we investigated stomatal traits (SD, SR, GCL, g_s_) in seven spring wheat genotypes under field conditions to assess their genetic and environmental variation, examine their relationships with grain yield, and provide insights for future breeding strategies. Agronomic traits of the studied genotypes have been reported elsewhere (Ingver *et al*., 2024) along with the effects of two nitrogen regimes used: N150, the typical fertilization rate for spring wheat in the Baltics and N75, representing a less intensive, environmentally friendly management approach (Jansone *et al*., 2024). Fully expanded flag leaves of spring wheat were measured with porometer for abaxial and adaxial g_s_ values and then collected for further stomatal anatomical analyses in 2022 and 2023. The vegetative period of 2022 was fairly average in terms of weather, with a slightly warmer, sunnier and drier June than the long period average (Fig. S1). In contrast, 2023 was characterized by a very dry and sunny May-June, followed by an extremely rainy July (Fig. S1). Likely due to these weather differences, grain yields in 2022 ranged between 5200-6900 kg ha^-1^, whereas in 2023 they were smaller (2800-4600 kg ha^-1^).

Genotype and year had significant main effects on SD and GCL values of adaxial and abaxial leaf surfaces, but only genotype affected adaxial to abaxial stomatal ratio (SR) (Table 1), suggesting that SR is largely genetically determined and less sensitive to environmental conditions than SD and GCL. Nitrogen fertilization had no significant effect on stomatal traits except abaxial GCL, but it increased flag leaf area and had a small positive effect on grain yield. In 2023, most of the cultivars showed significantly higher adaxial and abaxial SDs than in 2022 (Fig. 1a-b). Higher SDs in 2023 were associated with significantly smaller leaf areas due to early season drought (Fig. 1c). Leaf area has been found to correlate with SD, either negatively or positively (Salisbury, 1928; Brodribb *et al*., 2013; Peel *et al*., 2017; Wall *et al*., 2023). Stomata are formed early in leaf development before the full leaf expansion is reached (Gay & Hurd, 1975; Hepworth *et al*., 2018; Nunes *et al*., 2020). Grass stomatal development follows a strict pattern and adjusting SD to the environment may happen via changes in stomatal spacing within stomatal files or between them, or by changes in the number of stomatal files (Stebbins & Shah, 1960; McKown & Bergmann, 2020). Different abiotic stresses suppress leaf expansion growth, which relies on cell turgor and is strongly affected by leaf water status, leading to smaller leaves under stress conditions (Munns *et al*., 2000; Munns & Tester, 2008; Driesen *et al*., 2020). In 2023, drought started to develop already in May and intensified in June, hence, both stomatal development and expansion of flag leaves were affected by it. Drought-induced suppression of cell expansion resulted in about 2.5 times (average of all genotypes) smaller flag leaf area in 2023 than in 2022 (Fig. 1c).

**Table 1.**
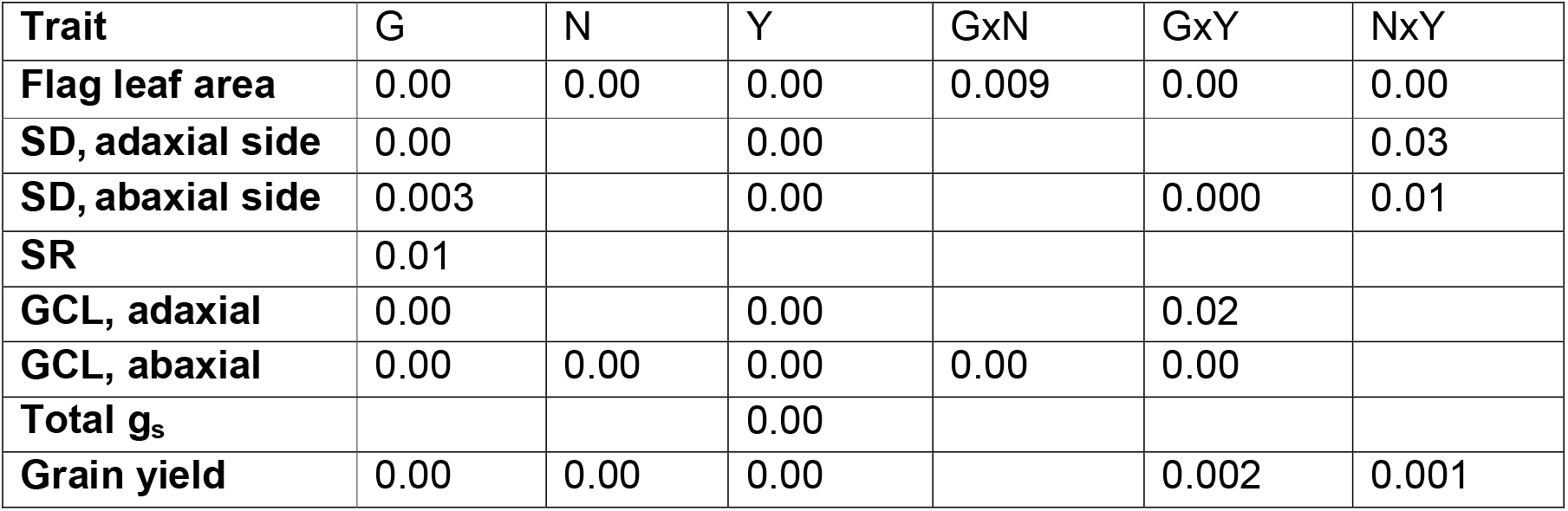
Results of ANOVA (p values) for the effects of genotype (G), nitrogen treatment (N), and year (Y) on studied flag leaves and their stomatal traits of 7 wheat genotypes. Empty cells indicate no significant effect, GxYxN interaction was never significant.

**Figure 1.**
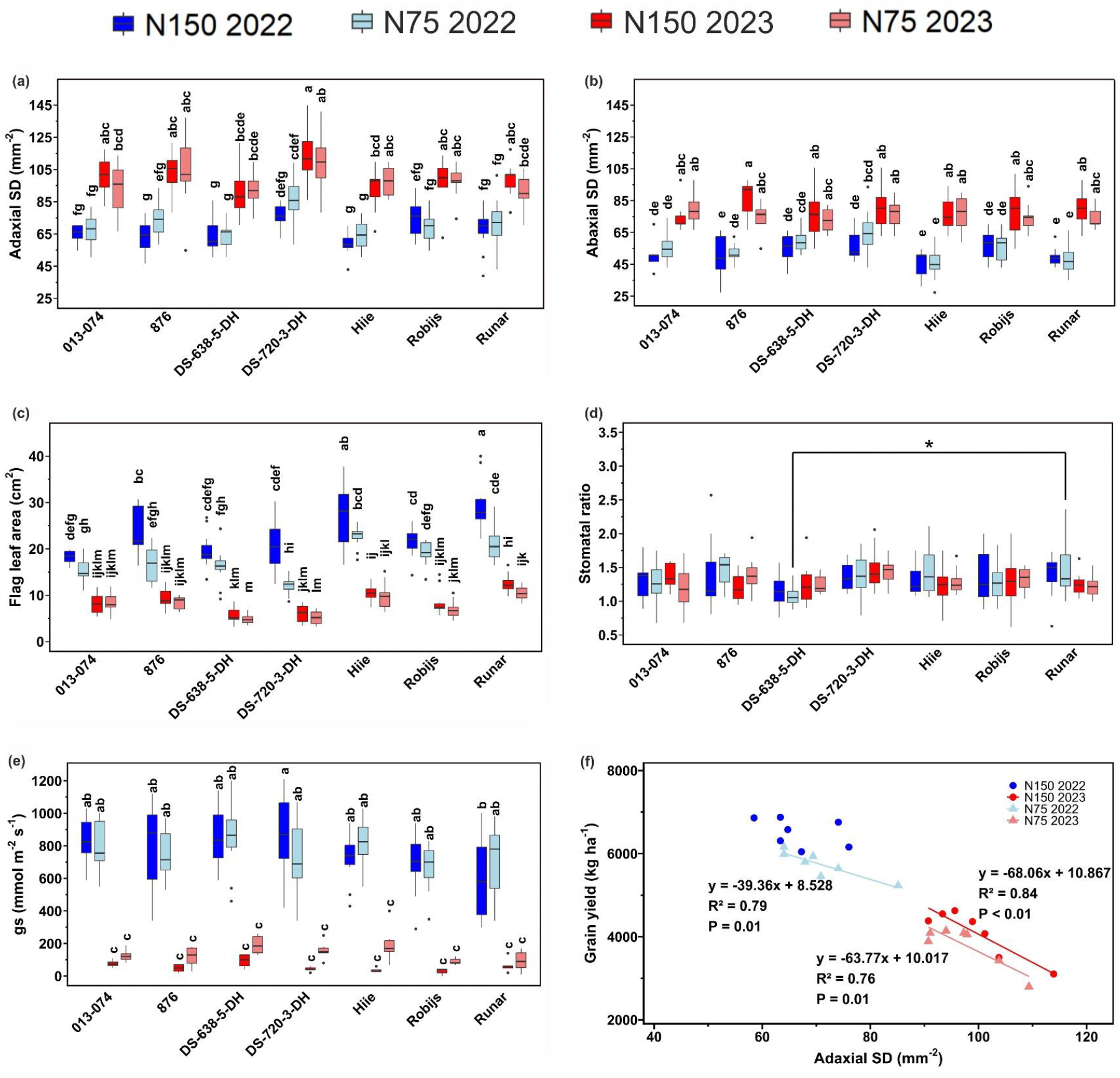
Stomatal traits of wheat flag leaves. a) Adaxial SD (stomatal density), n=10-12. b) Abaxial SD, n=10-12. c) Flag leaf area, n=12. d) Stomatal ratio calculated as a ratio of adaxial stomatal density to abaxial stomatal density. e) Total stomatal conductance calculated as sum of adaxial and abaxial stomatal conductances, n=5-12. f) Relationship between adaxial stomatal density and grain yield in 2022 and 2023, with regression lines and statistical significance (R^2^ and P-values). Box plots in panels (a–e) represent the distribution of values, with the boxes spanning the interquartile range (IQR) and the median indicated by a horizontal line. The whiskers extend to the non-outlier range, and potential outliers are shown as solid dots. The colours represent nitrogen treatments and years: blue for N150 (2022), light blue for N75 (2022), red for N150 (2023), and light coral for N75 (2023). Different letters for panels (a-c, e) and asterisk above the bracket for panel d denote statistically significant differences (GLM with Tukey *post hoc* test) between the genotypes.

In 2022, variation in flag leaf adaxial and abaxial SDs between genotypes was 1.46 and 1.42 times, respectively, whereas variation was less, 1.26 and 1.19 times, in the unfavourable year of 2023 (Fig. 1a-b). These values indicate that SD provides a valuable source of genetic diversity that has not yet been harnessed in breeding. As previously found in wheat (Wall *et al*., 2023), more stomata were detected on the adaxial than abaxial leaf surface as the values of SR were always above 1 (Fig. 1d). In 2022, guard cells were longer than in 2023 and adaxial guard cells were always longer than corresponding abaxial guard cells (Fig. S2a-b). Previously detected significant negative relationship between SD and GCL (Franks & Beerling, 2009; Haworth *et al*., 2023), was found only when years were pooled. Flag leaf total stomatal conductance of all genotypes was significantly lower in 2023 compared with 2022 due to drought (Fig. 1e). In 2022, g_s_ was similarly high in all genotypes.

Next, we analysed the relationships between stomatal traits and wheat grain yield by fitting linear regression models (Table S1). Interestingly, a strong significant negative relationship between adaxial SD and grain yield was detected for N75 plots in 2022 and 2023 and for N150 plots in 2023. (Fig. 1f). At the same time, abaxial SD was not significantly related with yield (Table S1). Neither total g_s_, nor g_s_ of leaf sides separately was related with grain yield in individual years (Table S1). This suggests that the significant negative relationship between adaxial SD and grain yield was not due to higher water loss of genotypes with higher adaxial SD, as g_s_ combines SD and stomatal aperture width to determine the diffusion of water vapour out of the leaf. This point is supported by a similarly strong decrease in g_s_ values of all wheat genotypes in response to water stress of 2023 compared with 2022, showing that all genotypes closed their stomata under drought (Fig. 1e). Alternatively, single point measurements of g_s_ taken around midday did not reveal the differences in diurnal water loss between genotypes. For example, genotypic variation in nocturnal stomatal conductance has been detected for wheat (McAusland *et al*., 2021); and higher adaxial SD could result in larger night-time water loss especially in Northern countries with dusky rather than dark summer nights.

Adaxial SD was the highest in the genotype DS-720-3-DH in both years, whereas the genotypes with the lowest adaxial SDs were Hiie and DS-638-5-DH (Fig. 1a). In 2021-2022, the phenotypic diversity of 300 spring wheat genotypes from the Nordic-Baltic region was studied and DS-720-3-DH ranked among superior genotypes due to its high grain yield, grain protein content, test weight, and thousand kernel weight (Jansone *et al*., 2024; Ingver *et al*., 2024). Our study indicates that because of its high adaxial SD, DS-720-3-DH is not the best parental material in breeding for stressful conditions as also supported by its very low yield in the drought year of 2023. All stomatal traits were significantly related with grain yield within the pooled dataset, except SR (which was negatively related with grain yield in N75 plots in 2023; Table S1). However, for pooled data, these significant relationships are rather due to environmental differences between the years and not due to differences between genotypes.

In conclusion, the considerable variation in adaxial SD between wheat genotypes in individual years and between different years was highly indicative of yield: the less stomata on the adaxial side, the larger the grain yield. This result was found for 2023 characterized by serious early season drought and for N75 plots in 2022. At this point, we can only hypothesize about the reasons behind the negative relationship between adaxial SD and grain yield: as Junes of both years were warmer, sunnier, and drier than usual, larger dehydration associated with more adaxial stomata seems the most probable cause. Different pathogen sensitivity due to variation in adaxial SD is less probable in these conditions, but cannot be excluded. For N150 plots in 2022, there was no significant relationship between adaxial SD and grain yield. Possibly, under favourable environmental and fertilization conditions, such as those that led to the highest yield detected during this experiment in N150 plots in 2022 (6512 ± 129 kg ha^-1^ as an average of all genotypes), the negative effect of higher adaxial SD is not manifesting itself, but does so under stress conditions. Correlations between SD, net assimilation rate, and yield have been detected in previous studies (Gaskell & Pearce, 1983; McAusland *et al*., 2021; Kumar *et al*., 2021). In rice, drought-resistant lines showed consistently lower adaxial stomatal densities compared with the drought-susceptible ones and adaxial SD negatively correlated with yield under high vapour pressure deficit (Kumar *et al*., 2021). In maize, hybrids showing higher carbon exchange rates had larger adaxial SD and lower stomatal resistance (Gaskell & Pearce, 1983). In wheat, higher abaxial, but not adaxial SD led to greater numbers of heavier seed (McAusland *et al*., 2021), whereas in other studies, neither abaxial SD of wheat flag leaves (Liao *et al*., 2005) nor adaxial SD of wheat flag leaves (Shahinnia *et al*., 2016) correlated with grain yield. Our study simultaneously measured and distinguished between leaf surfaces to highlight their distinct effects on yield. Stomatal distribution has been recognized as a functional trait for improved photosynthesis (Lawson & Blatt, 2014; Xiong & Flexas, 2020); however, higher-than-optimal adaxial SD values may result in yield losses of wheat. Its reasons and whether the same is true for crops with lower SR values await further studies.

## Materials and methods

### Planting material and field experiments

Seven spring wheat (*Triticum aestivum* L.) genotypes representing breeding lines and released varieties developed in the spring wheat breeding programs of the Baltic States and Norway were studied. The field experiments were carried out at the Centre of Estonian Rural Research and Knowledge (METK, Estonia), Estonia (58°45’N, 26°24’E) in 2022–2023. The wheat genotypes were sown (plot size of 10 m^2^, and row spacings of 12.5 cm) under two nitrogen (N) fertilization levels applied before sowing: 75 kg ha^-1^ N (referred to as N75) and 150 kg ha^-1^ (referred to as N150). Two plots per genotype and N level were sown on 03.05.2022 and 25.04.2023. At maturity, the plots were harvested mechanically (dates 18-25.08.2022 and 14-21.08.2023), and the grain yield (kg ha^-1^) was expressed on a 14% grain moisture basis.

### Stomatal conductance (g_s_), stomatal density (SD), and leaf area measurements

The timing of flag leaf measurements was chosen to study fully expanded leaves at their maximum physiological activity. Adaxial and abaxial leaf sides of six flag leaves per plot were measured with a LI-600 (LI-COR) hand-held porometer to get stomatal conductance, g_s_. In further data analysis, total g_s_ was calculated as the sum of adaxial and abaxial stomatal conductances. The measured leaves were then harvested and taken to the laboratory for leaf area and stomatal density measurements. Leaves were laid flat on a table, photographed, and their projected areas were determined with ImageJ software (Schneider *et al*., 2012). For SD measurements, the central area of the flag leaf was sampled, and impressions of both the adaxial and abaxial leaf surfaces were made using dental silicone. Specifically, Speedex light body (Coltene/Whaledent AG, Alstätte, Switzerland) was used for the adaxial side and oranwash L (Zhermack, www.zhermack.com) for the abaxial side. Secondary imprints made with nail varnish were collected from the silicone and were transferred onto microscope slides with the help of transparent tape. An area of 0.26 mm^2^ from each imprint was photographed under a microscope (Kern OBF 133; Kern & Sohn GmbH; www.kern-sohn.com) at 200x magnification. Stomatal density, SD (stomata per mm^2^), and guard cell length, GCL, were determined from these images using ImageJ software (Schneider *et al*., 2012) and stomatal ratio calculated as the ratio of adaxial SD to abaxial SD.

## Statistical analyses

Analysis of variance (GLM procedure) was used to evaluate the main effects of genotype, N fertilization and year on leaf area, stomatal traits and grain yield (Statistica, version 7.0, StatSoft Inc., Tulsa, OK, USA). Comparisons between measured variables were performed using 3-way analysis of variance (ANOVA) followed by Tukey post hoc test to compare the individual means with the R scripting language, version 4.4.2. Simple linear regression analysis was performed to detect significant relationships between studied traits and grain yield. All effects were considered significant at p < 0.05.

## Acknowledgements

This study was funded by the Estonian Research Council, PSG404 and PRG1620, and project “NOBALwheat – breeding toolbox for sustainable food system of the NOrdic BALtic region”, Research Council of Lithuania (LMTLT) contract No. S-BMT21–3 (LT08-2-LMT-K-01–032). We are grateful to Mikk Välbe, Isabella Siil, Daana Morozova, Marilin Poogen, Katarína Geletková and Dmitrii Kliaus for their help with sample collection, stomatal imaging and counting. The research was conducted using the TAIM Plant Biology Infrastructure funded by the Estonian Research Council (TT5).

## Author contributions

H.H initiated the studies concerning stomatal imaging of different leaf surfaces; H.H. and E.M. conceived and designed the study; G.B. and M.B. were involved in the generation of plant lines; K.S, P.J., E.I., I.Tulva, B.K-D., A.I., I.Tamm., H.H., and E.M. collected and analysed data, K.S. and E.M. wrote the manuscript with input from all authors.

