## Supplementary material for "Higher adaxial stomatal density is associated with lower grain yield in spring wheat": Table S1

**Supplemental Table 1.** Significant linear relationships between dependent variable (grain yield) and different leaf and stomatal traits (flag leaf area; SD, ad: adaxial stomatal density; SD, ab: abaxial stomatal density; SR: adaxial to abaxial stomatal ratio; g_s_, Total: stomatal conductance of adaxial and abaxial leaf sides summed; g_s_, ad: adaxial stomatal conductance; g_s_, ab: abaxial stomatal conductance) are presented for individual years and for a pooled dataset across 2022-2023. The correlation coefficients (R values) for significant relationships (p<0.05) are shown. Empty cells indicate no significant relationship.

| **Traits** | **2022, N75** | **2022, N150** | **2023, N75** | **2023, N150** | **2022-23**  **N75** | **2022-23**  **N150** |
| --- | --- | --- | --- | --- | --- | --- |
| **Flag leaf area** |  |  |  |  | 0.89 | 0.87 |
| **SD, ad** | -0.89 |  | -0.87 | -0.92 | -0.97 | -0.97 |
| **SD, ab** |  |  |  |  | -0.88 | -0.93 |
| **SR** |  |  | -0.84 |  |  |  |
| **GCL, ad** |  |  |  |  | 0.64 | 0.56 |
| **GCL, ab** |  |  |  |  | 0.61 | 0.75 |
| **g_s_, Total** |  |  |  |  | 0.93 | 0.93 |
| **g_s_, ad** |  |  |  |  | 0.92 | 0.93 |
| **g_s_, ab** |  |  |  |  | 0.91 | 0.93 |
