## Supplementary material for "Higher adaxial stomatal density is associated with lower grain yield in spring wheat": Fig. S1

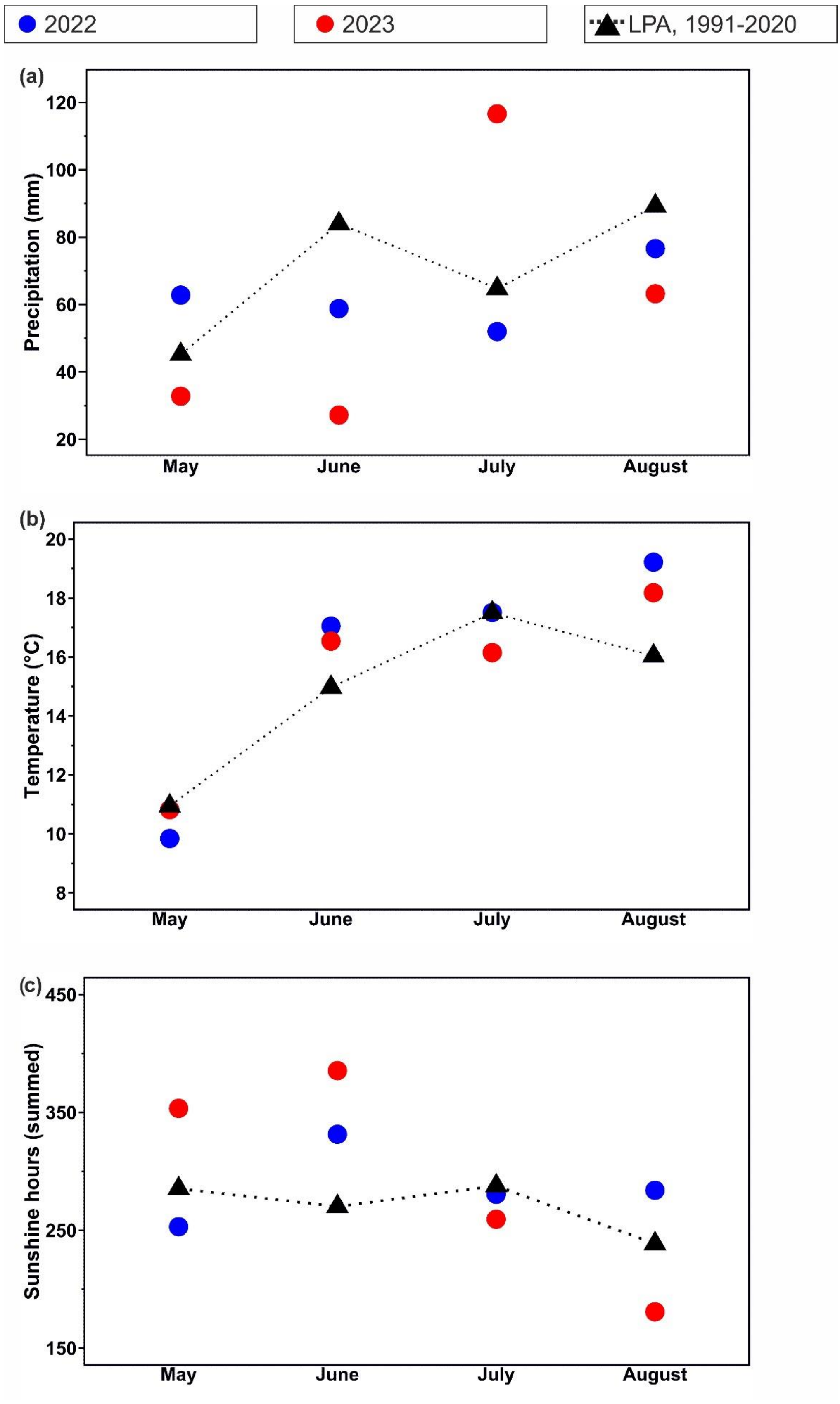


Figure S1. Weather data for experimental years: (a) Sum of precipitation, (b) Average temperature, and (c) Sum of sunshine hours for May-August, 2022-2023 and respective LPA (long period averages) (1991-2020) in Jõgeva experimental station.
