## Supplementary material for "Higher adaxial stomatal density is associated with lower grain yield in spring wheat": Fig. S2

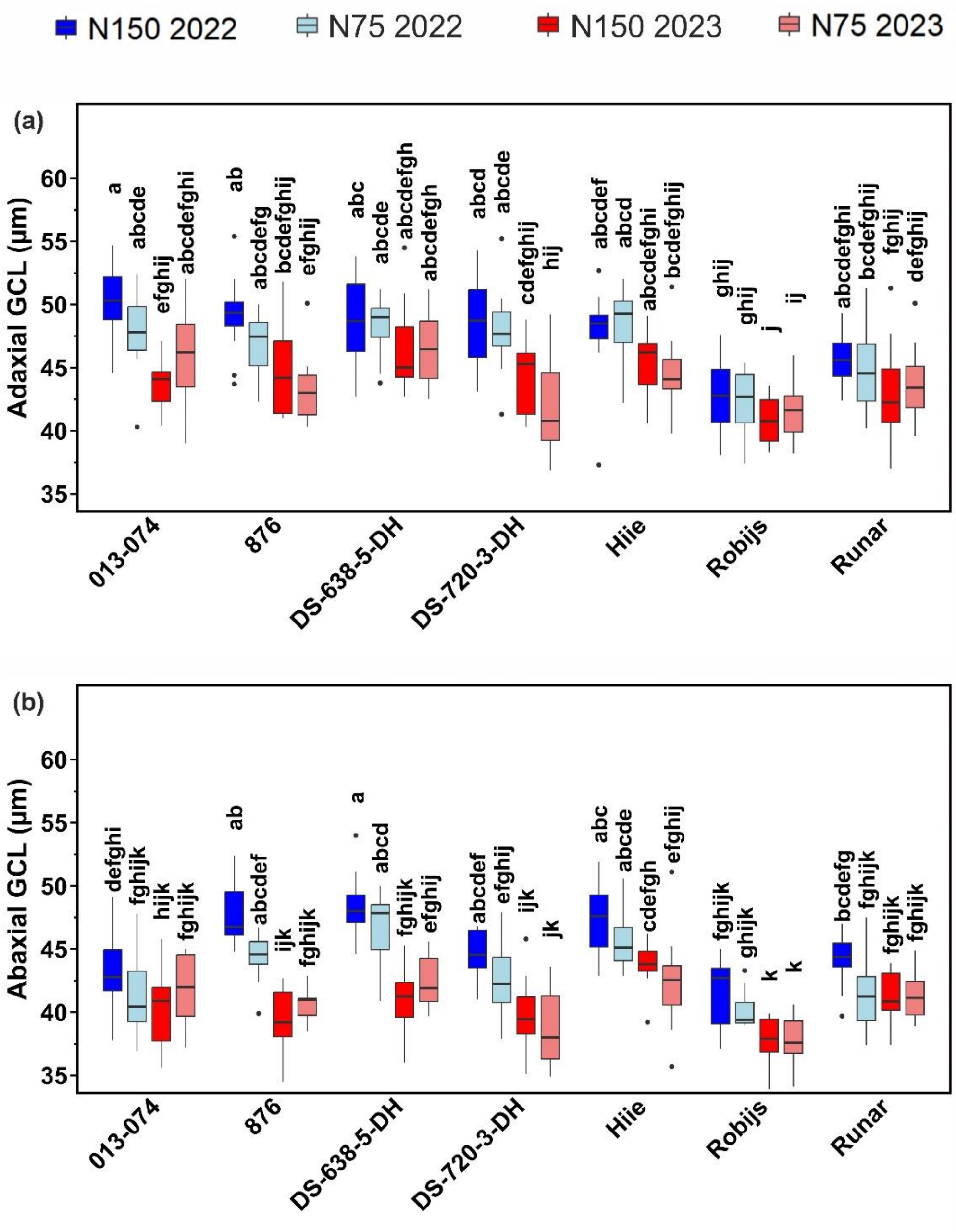


**Figure S2. Guard cell lengths (GCL) of adaxial and abaxial leaf surfaces.** (a) Adaxial GCL, n=10-12. (b) Abaxial GCL, n=10-12. Box plots illustrate the distribution of values, with the boxes representing the interquartile range (IQR) and the median marked by a horizontal line. The whiskers extend to the range excluding outliers, while potential outliers are displayed as solid dots. Colours indicate nitrogen treatments and years: blue for N150 (2022), light blue for N75 (2022), red for N150 (2023), and light coral for N75 (2023). Statistically significant differences are denoted by different letters (GLM with Tukey *post hoc* test).
